# Induction of long-lived potential aestivation states in laboratory *An. gambiae* mosquitoes

**DOI:** 10.1101/2020.04.14.031799

**Authors:** Benjamin J. Krajacich, Margery Sullivan, Roy Faiman, Laura Veru, Leland Graber, Tovi Lehmann

**Affiliations:** Laboratory of Malaria and Vector Research, National Institute of Allergy and Infectious Diseases, National Institutes of Health, Rockville, MD, 20852

**Keywords:** aestivation, dry season, *Anopheles*, malaria

## Abstract

**Background:** How *Anopheline* mosquitoes persist through the long dry season in Africa remains a gap in our understanding of these malaria vectors. To span this period in locations such as the Sahelian zone of Mali, mosquitoes must either migrate to areas of permanent water, recolonize areas as they again become favorable, or survive in harsh conditions including high temperatures, low humidity, and an absence of surface water (required for breeding). Adult mosquitoes surviving through this season must dramatically extend their typical lifespan (averaging 2-3 weeks) to 7 months. Previous work has found evidence that the malaria mosquito *An. coluzzii*, survives over 200 days in the wild between rainy seasons in a presumed state of aestivation (hibernation), but this state has so far not been replicated in laboratory conditions. The inability to recapitulate aestivation in the lab hinders addressing key questions such as how this state is induced, how it affects malaria vector competence, and its impact on disease transmission.

**Methods:** We compared survivorship of mosquitoes in climate-controlled incubators that adjusted humidity (40-85% RH), temperature (18-27 °C), and light conditions (8-12 hours of light). A range of conditions were chosen to mimic the late rainy and dry seasons as well as relevant extremes these mosquitoes may experience during aestivation.

**Results:** We found that by priming mosquitoes in conditions simulating the late wet season in Mali, and maintaining mosquitoes in reduced light/temperature, mean mosquito survival increased from 18.34 +/− 0.65 to 48.02 +/− 2.87 days, median survival increased from 19 (95% CI: 17-21) to 50 days (95% CI: 40-58), and the maximum longevity increased from 38 to 109 days (*p*-adj < 0.001). While this increase falls short of the 200+ day survival seen in field mosquitoes, this extension is substantially higher than previously found through environmental or dietary modulation and is hard to reconcile with states other than aestivation. Future work will expand on these findings, looking to further extend the gains in life span while also investigating transcriptional changes in genes affecting aging, metabolism, and epigenetic regulation; hormone and nutrient levels; and vector competence for malaria causing parasites.

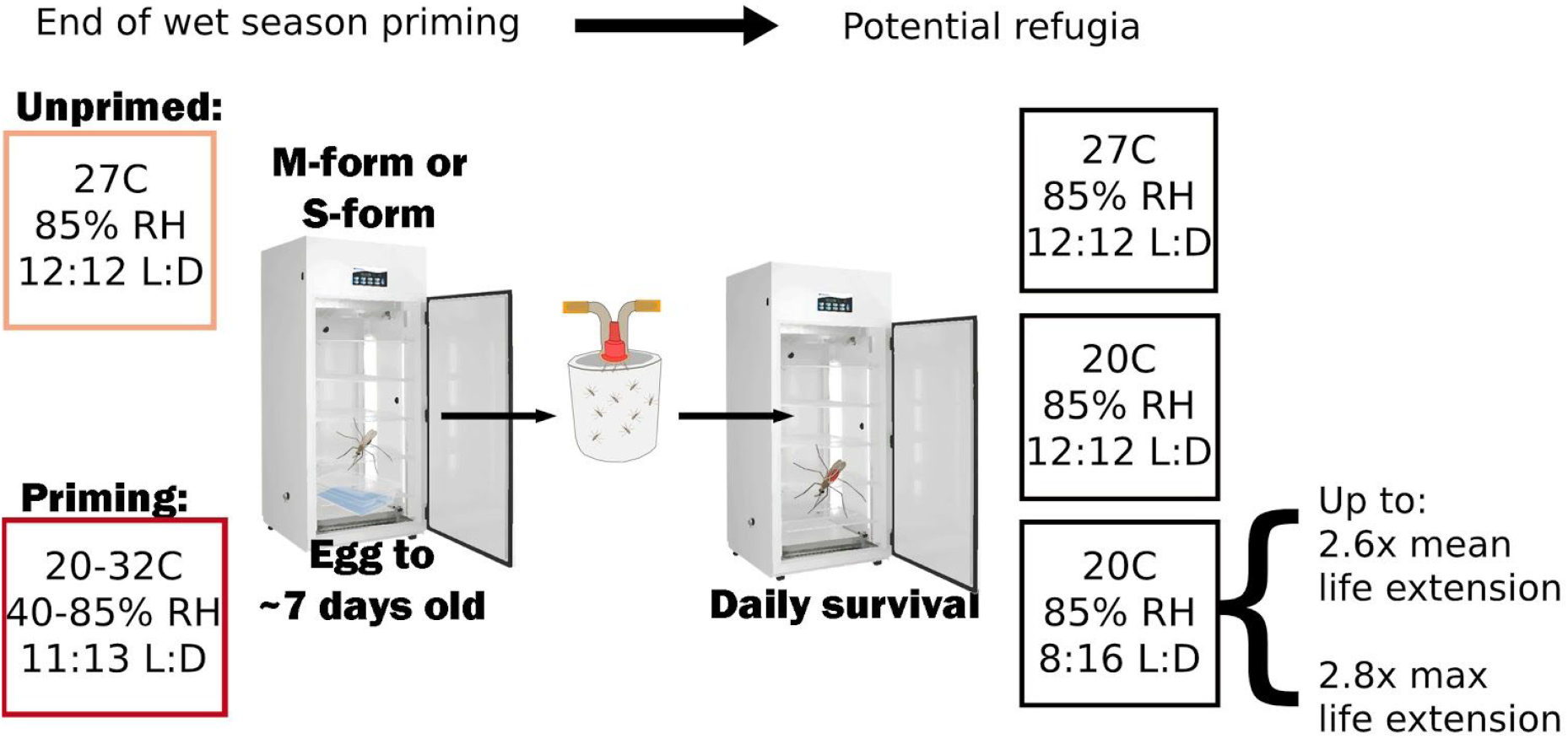

## Background

The goal to end the transmission of malaria causing *Plasmodium* spp. parasites has achieved significant gains since the recent peak of the early 2000s, though it appears progress may have slowed in sub-Saharan Africa in the last two years [1]. Currently, the majority of infected individuals (>90%) are living in meso or hypoendemic (defined as a *Plasmodium falciparum* parasitaemia rate in children aged 2-10 between 1-50%) areas with seasonal transmission [2]. While Mali has a strong seasonal transmission cycle, with ~7 months of the year largely unconducive to mosquito presence, the rates of malaria have remained consistent [3], even with increasing intervention in the form of bed nets [4], insecticide residual spraying [5], and other interventions. The reasons for this disease persistence are multifactored, but it is likely that the persistent nature of the vectors themselves play a role. Insecticide resistance has emerged broadly to all utilized insecticides [6], eradication of *An. gambiae* has only been possible in a few areas of the globe [7, 8], and the resilient seasonal return of malaria-causing vectors has been long known though with poorly understood mechanisms.

Evidence for long-lived aestivation, or dry-season diapause phenotypes have been described for over 100 years in *Anopheles* mosquitoes [9]. In Burkina Faso, collections of larvae and adult *An. gambiae* from the area around Bobo-Dioulasso showed the ability to live greater than 150 days when held in houses that were comparatively cooler, more humid, and less windy than the ambient air [10]. A similar finding was also reported with *Anopheles gambiae* (likely contemporary *An. arabiensis*) from Sudan, where in the absence of suitable breeding habitat, mosquitoes were found resting in “dwelling huts in dark places between the thatched roofs and the median longitudinal beams” during the entirety of the 9 month dry season [11]. More contemporary work in Mali has highlighted the unique patterns in seasonal abundance of the three sibling species of the *An. gambiae* s.l. complex, with *An. coluzzii* mosquitoes showing presence throughout the arid 7 month dry season, *An. gambiae* s.s. being absent, and *An. arabiensis* showing an intermediate phenotype [12, 13]. The *An. coluzzii* mosquitoes that do appear during the late dry season peak (a yearly, high density, 1-2 week event), host-seek normally, but exhibit reproductive depression compared with the wet season mosquitoes [14]. However, while this phenotype has been repeatedly described over the last century, the methodology to induce such a long-lived state with laboratory, colony mosquitoes has remained beyond reach.

Laboratory models of overwintering diapause have been reliably induced in other *Culicidae* such as *Culex pipiens* and *Aedes albopictus* in response to relatively simple approximations of winter conditions of reduced temperature and photoperiod [15, 16]. However, multiple studies performed to induce aestivation via oviposition deprivation [17], adult photoperiod and temperature modulation [18], and adult dietary restriction [19], have failed to produce the dramatic lifespan extension typical of aestivation with only a modest increase in median lifespan from 20 to 30 days and maximum life spans of up to 64 days. This study aims to build upon past cues tested for adult aestivation induction. This includes more consideration of temperature and photoperiod changes that may induce or prime mosquito larvae and early adults for aestivation during the late wet season period, as well as temperature and photoperiod cues that would be more consistent with refugia conditions of the early dry season that could be important for the maintenance of the state. We evaluate longevity as a primary outcome and describe some morphological and developmental characteristics surrounding these conditions.

## Materials and methods

### Mosquito rearing

Two strains of *An. gambiae* s.l. complex mosquitoes were utilized in this experiment. The first was the Thierola strain of *An. coluzzii* that were founded as a colony from six wild-caught females from Thierola, Mali (13.6586, −7.21471) in November, 2012 [18]. The second was the N’dokayo strain of *An. gambiae* s.s. (MRA-1278) provided through the Malaria Research and Reference Reagent Resource Center (MR4), originally colonized in 2008 from Cameroon (5°30.723’N 14°07.497’E) [20]. All mosquito larvae were reared in plastic trays (30 × 25 × 7cm) with 1.5 L of dechlorinated water. In the first 24 hours after emergence, trays were provided with 5 mL of a 4% w/v baker’s yeast slurry [21], and subsequently provided with finely-ground TetraMin fish food (~ 0.1g daily, Tetra Inc., Melle, Germany) [19]. Pupae were picked daily and kept in separate adult cages with sugar provided *ad libitum* as 10% karo dark syrup on cotton balls refreshed daily. Normal insectary conditions were 27 °C, 85% humidity, and 12:12 hours of light:dark with 30 minutes of sunrise/sunset period. All blood feeds were performed using glass, water-jacketed blood feeders, with 1 mL of human blood (Interstate Blood Bank, Inc.) hooked to a recirculating water bath kept at 37 °C.

### Experimental groups

To evaluate possible conditions relevant for aestivation induction, we utilized “priming” conditions for mosquitoes based on the downturn in numbers of *An. coluzzii* mosquitoes at the end of September/October [12]. Using data from a weather monitoring station in Thierola, Mali, we calculated mean daily temperature and humidity profiles for each month and used a climate controlled incubator (I-36VL, Percival Scientific, Perry, IA, USA) to approximate conditions during this time. This temperature and humidity profile alternates between night conditions of 20 °C and 85% RH and day conditions of 32 °C with 40% RH and a 11:13 light:dark cycle (Additional File 1).

**Additional File 1:**
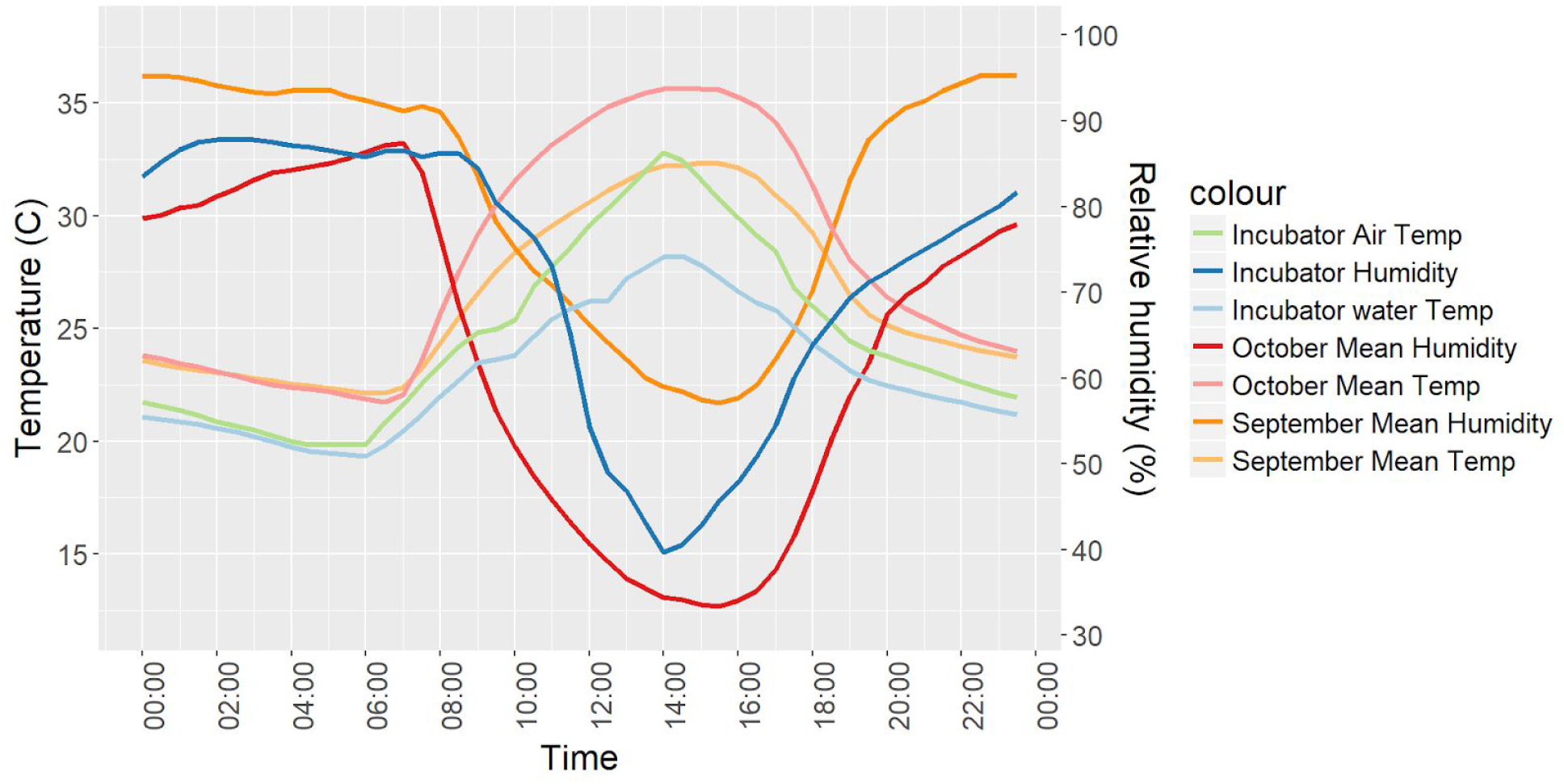
Temperature and Humidity profiles for Thierola, Mali (Red, Orange) and the priming incubator (Green, Blue) as recorded by a HOBO temperature and humidity logger.

To determine initial conditions relevant for aestivation maintenance, a range of refugia-like conditions were chosen based on values to the extreme end of what is likely to be experienced during the Sahelian dry season. In the first experiment, mosquitoes were kept in a climate controlled incubators with initial refugia conditions set to either 18 °C or 22 °C, 8:16 light:dark hours of broad spectrum led light (Transcend T8 13.5 and 22W bulbs, Hort Americas, Bedford, TX), and 85% humidity, with a group left under the priming conditions and a group under standard insectary conditions. During the second experimental trial, the refugia conditions were 20 °C, 8:16 hours of broad spectrum led light, and 85% humidity. Subsequent experiments were limited to either 20 or 27 °C, with 8 or 12 hours of light and consistent 85% relative humidity.

### Survival analysis and statistics

To determine the effects from differing environmental conditions, priming, and mosquito species, non-parametric and parametric survival analyses were performed using the ‘survminer’ and ‘flexsurv’ packages in R using R-Studio (R version 3.6.0, [22–25]). Not all variables passed the proportional hazards test, so we used accelerated failure time models to determine the effect size of each variable on survival time. We selected the best-fitting model distribution by comparing Akaike’s Information Criterion (AIC) values of each survival fit including variables.

### Morphological characteristics

Upon death, individual mosquitoes were placed into a 1.7 ml tube half filled with desiccant (Silica-gel orange, Cat. No. 10087. Sigma-Aldrich, St. Louis, MO) topped with a small ball of cotton. Mosquitoes were kept at −20 °C until dissection. Wing lengths were taken as previously described [18], and wing areas were calculated by determining the convex hull of the fourteen wing landmarks in R, discarding any samples that were missing landmarks. Differences in lengths and areas were compared via Kruskal-Wallis test with Dunn’s correction for multiple testing with the ‘dunn.test’ package in R [26].

## Results

### Effects of refugia and priming conditions on longevity

In the first trial to identify potential refugia conditions, we tested the survival of blood fed *An. coluzzii* mosquitoes held at 18 and 27 °C. Mosquitoes reared as larvae at 27 °C/12hr L:D when maintained in these conditions after emergence and blood feeding had a median lifespan of 21 days, with a maximum of 34 days. Mosquitoes reared at 27 °C/12hr L:D until 24 hours after the first blood meal and then transferred to 18 °C with a 8:16 L:D photoperiod significantly extended the median lifespan to 41 days, with a maximum of 80 days (*n*=59/group, *p* < 0.001, Figure 1A). Mosquitoes reared under end of wet season conditions (20-32 °C daily), then transferred to 18 °C, 22 °C, 27 °C, or maintained under end of wet season priming conditions, had median lifespans of 50, 46, 26, and 27 days and maximum lifespans of 102, 89, 44, and 52 days, respectively. Median lifespans were significantly different between normal insectary conditions and 18 °C, 22 °C (adjusted *p*-values < 0.001, Figure 1B).

**Figure 1:**
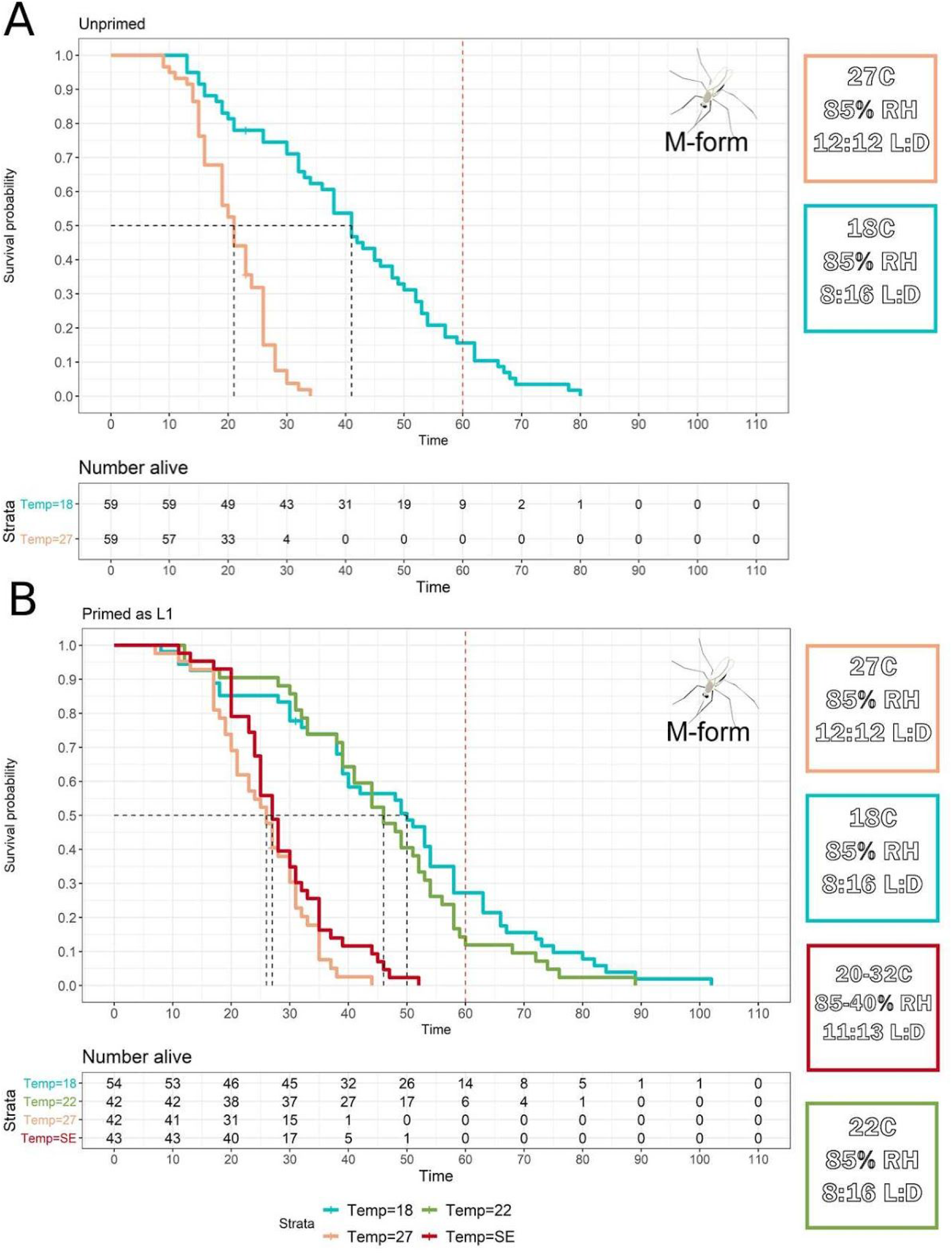
Kaplan-Meier Survival curves to evaluate priming and refugia conditions. **(A)** Bloodfed, unprimed *An. coluzzii* (M-form) mosquitoes reared in the listed conditions (18°C significantly higher survival, p < 0.001). **(B)** Bloodfed, primed from L1 in end of wet-season conditions, *An. coluzzii* mosquitoes reared in the listed conditions (18°C, 22°C significantly higher survival than 27°C and SE conditions p-adj < 0.001. 18°C vs 22°C and 27°C vs SE not significantly different, p-adj > 0.11)

### Effect of simulated induction and maintenance of aestivation on longevity’

Once the link between the refugia conditions and longevity had been established, we performed three additional survival experiments to determine the role of photoperiod, priming, species, and temperature on the longevity phenotype (Figure 2).

**Figure 2:**
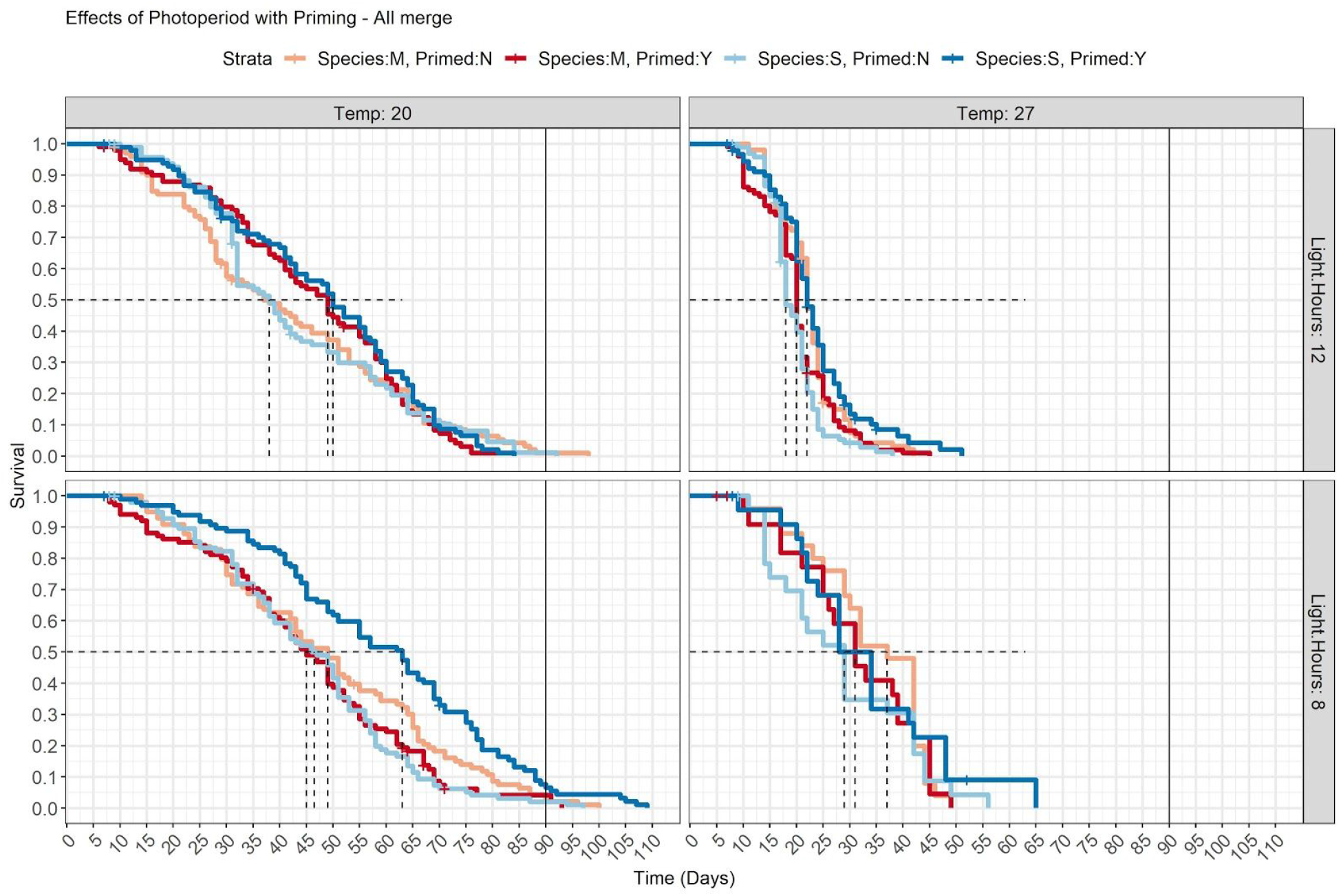
Kaplan-Meier Survival curves for each experimental group tested. The data from three independent trials is combined (*n*=100 mosquitoes/group), and presented for all groups except for 27 °C / 8:16 L:D cycle which was added during the last replicate (*n* = 25 mosquitoes/group). All 20°C significantly different from 27°C (adj p-val < 0.01). Priming significant for S-form mosquitoes at 20°C/8:16hr photoperiod (adj. p-val < 0.001)

Overall, as in the first experiments, we found significant extension of lifespan with both M and S form mosquitoes when maintained in cooler, darker environments than general insectary conditions with median survival ranging from 18 days (18-21 95% CI) in 27 °C / 12:12 hr L:D / unprimed / S-form mosquitoes to a maximum median survival of 63 days (55-69 day 95% CI) in 20 °C / 8:16 hr L:D / primed / S-form mosquitoes (Table 1). Priming had variable results depending on condition and species, i.e. at 20 °C with 12:12 L:D cycle, priming increased longevity for both M and S mosquitoes (median survival from 38 to 49 days and from 38 to 50 days, respectively), whereas at 20 °C with 8:16 L:D cycle, only S-form mosquitoes increased survival with priming (median survival from 46.5 to 63 days with priming in S-form, 49 to 45 days in M-form with priming).

**Table 1:**
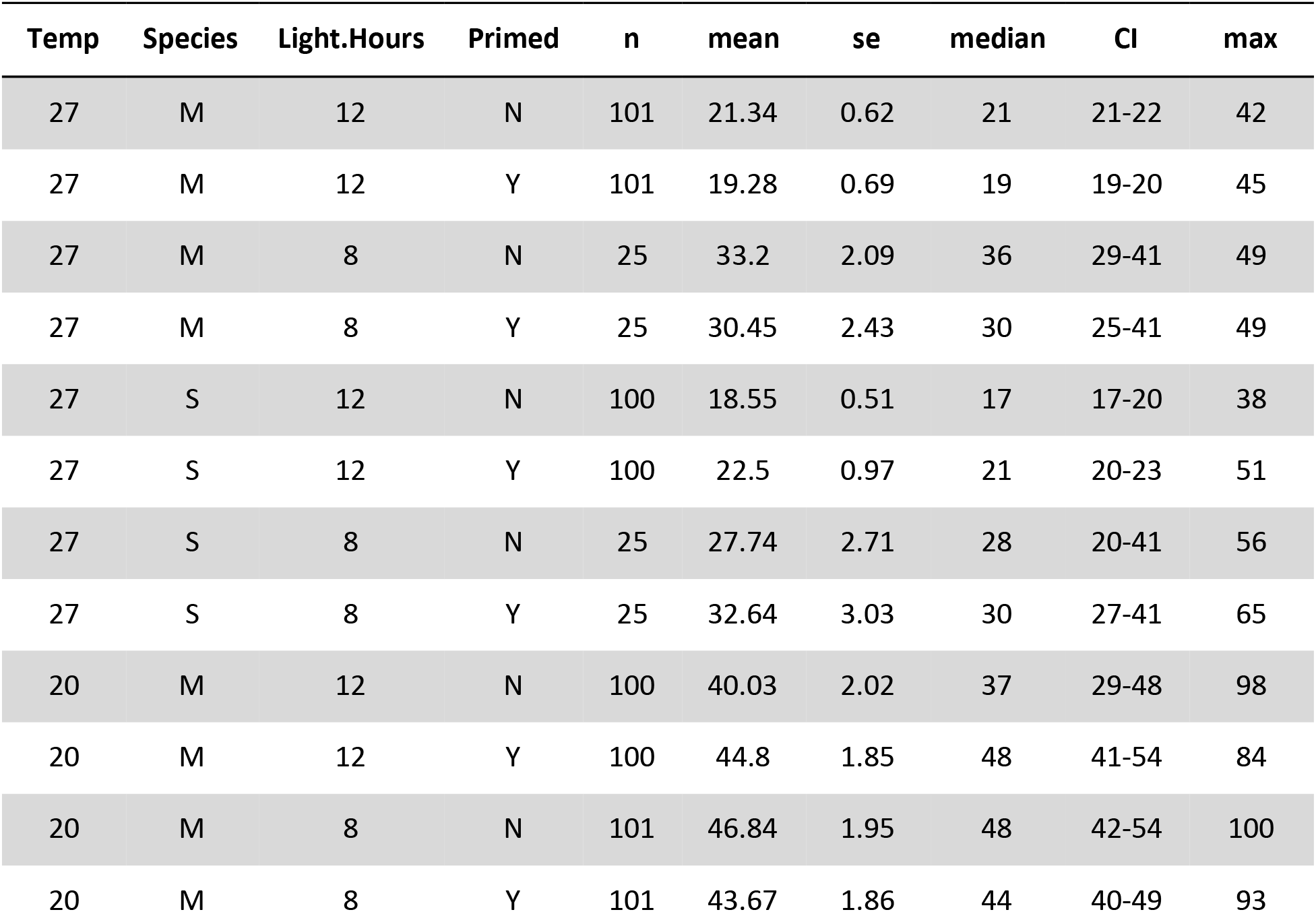
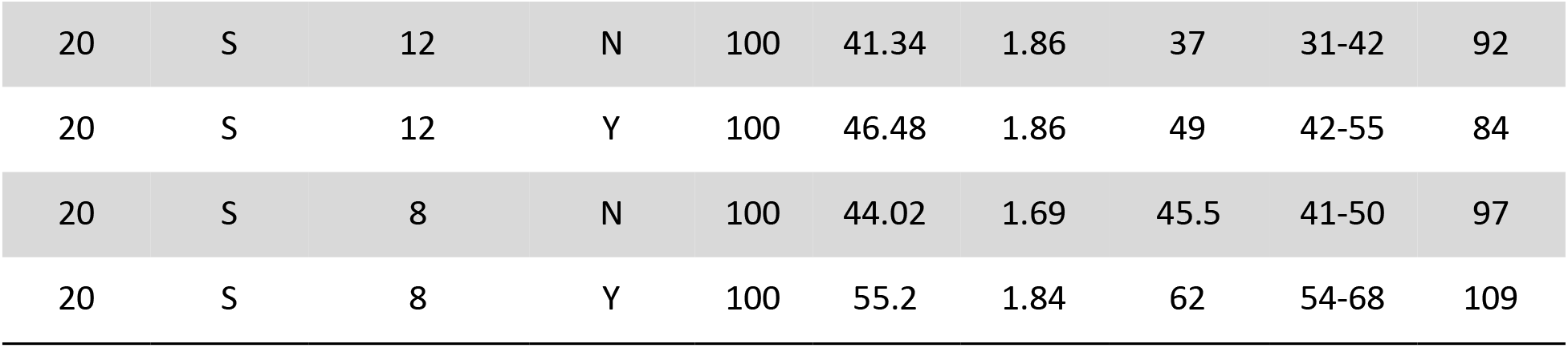
Conditions and survival characteristics for priming and temperature conditions. See also Additional File 2.

**Additional File 2:**
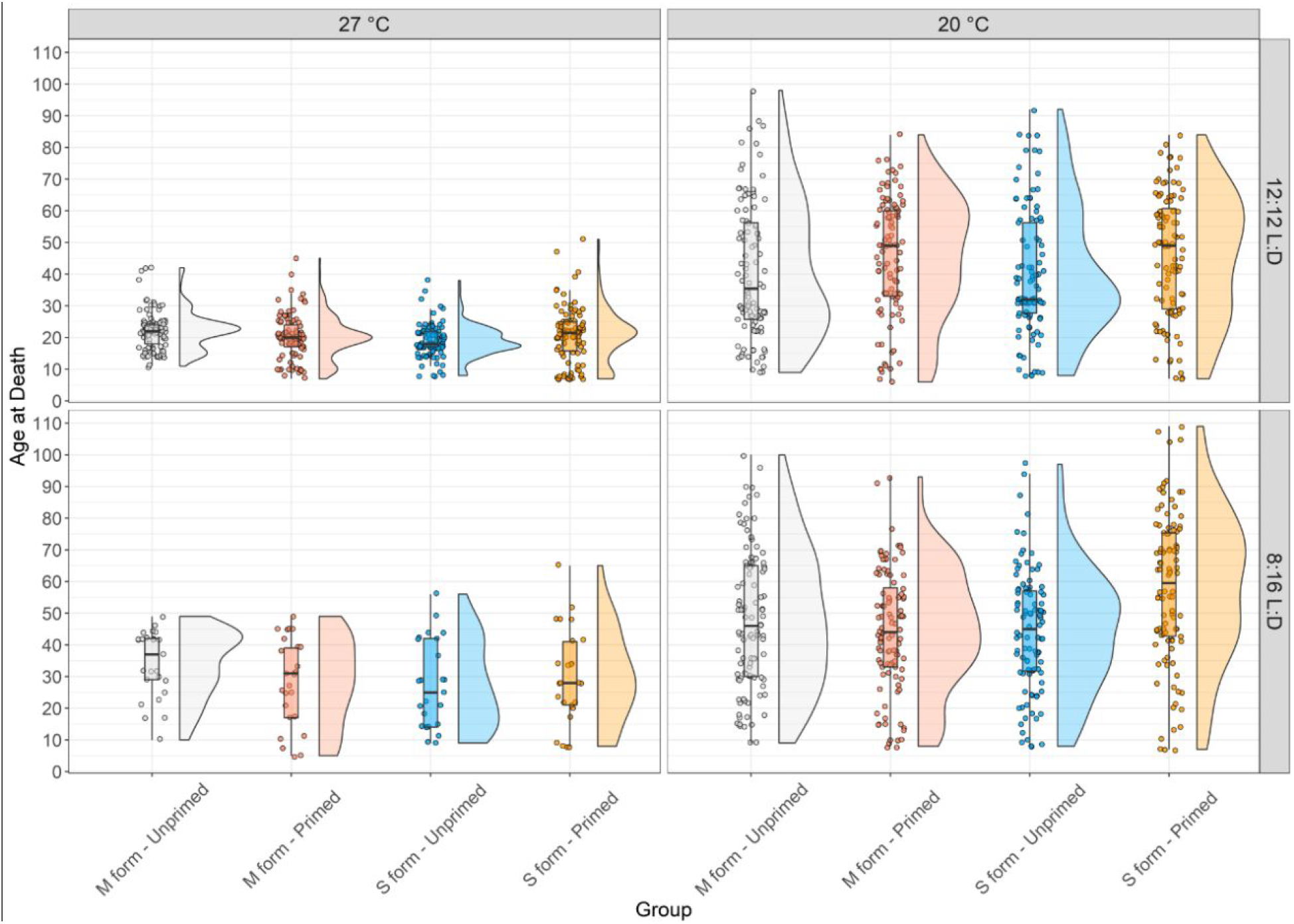
Boxplot and raincloud [27] plots showing the date of death ranges and distributions for each of the experimental groups.

To better understand the contribution of the treatment variables, we utilized an accelerated failure time model using a Weibull distribution, chosen as the best fit of our data based on AIC (Additional File 3). Using this model we evaluated the specific effects of each component via their event time ratio (time to death). We found that temperature, photoperiod, priming, and species significantly increased survival (*p*-values of <0.001, <0.001, <0.001, and 0.009, event time ratios of 2.02, 1.24, 1.19, 1.12, respectively, Table 2), indicating that storage at 20 °C vs 27 °C doubles survival, 8 hours of light a day vs 12 increased survival by 24%, end of wet season priming conditions over standard insectary increased survival by 19%, and M-form overall had slightly higher survival across all conditions by 12%. As we expected the species to differ in their response to these variables we allowed for an interaction term with species. These were largely insignificant, though there was an unexpected negative relationship with priming and M-form mosquitoes (ETR 0.81, p < 0.001)

**Table 2:** Significance of individual variables towards survival as estimated using an Accelerated Failure Time model with a Weibull distribution. Significant variables are bolded, and event time ratios (ETR) are presented. These can be interpreted as the time to death of the listed condition for that variable (i.e. a mosquito at Temp 20 °C has double the time to death of a mosquito at 27 °C). Interaction terms are designated with a colon between variables.

| variable | value | SE | z | p | ETR |
| --- | --- | --- | --- | --- | --- |
| (Intercept) | 3.107895 | 0.030706 | 101.2135 | 0 | 22.37 |
| **Temp (20C)** | **0.704108** | **0.031994** | **22.00753** | **2.4E-107** | **2.02** |
| **Light.Hours (8:16)** | **0.218615** | **0.0315** | **6.940067** | **3.92E-12** | **1.24** |
| **Primed (Yes)** | **0.1703** | **0.030607** | **5.564113** | **2.63E-08** | **1.19** |
| **Species (M)** | **0.109571** | **0.042466** | **2.580224** | **0.009874** | **1.12** |
| Temp20:SpeciesM | -0.02792 | 0.044161 | -0.63232 | 0.527178 | 0.97 |
| Light.Hours8:SpeciesM | -0.03256 | 0.043985 | -0.74027 | 0.459134 | 0.97 |
| **PrimedY:SpeciesM** | **-0.20976** | **0.042588** | **-4.92548** | **8.42E-07** | **0.81** |
| Log(scale) | -0.99224 | 0.022042 | -45.015 | 0 | 0.37 |

**Additional File 3:**
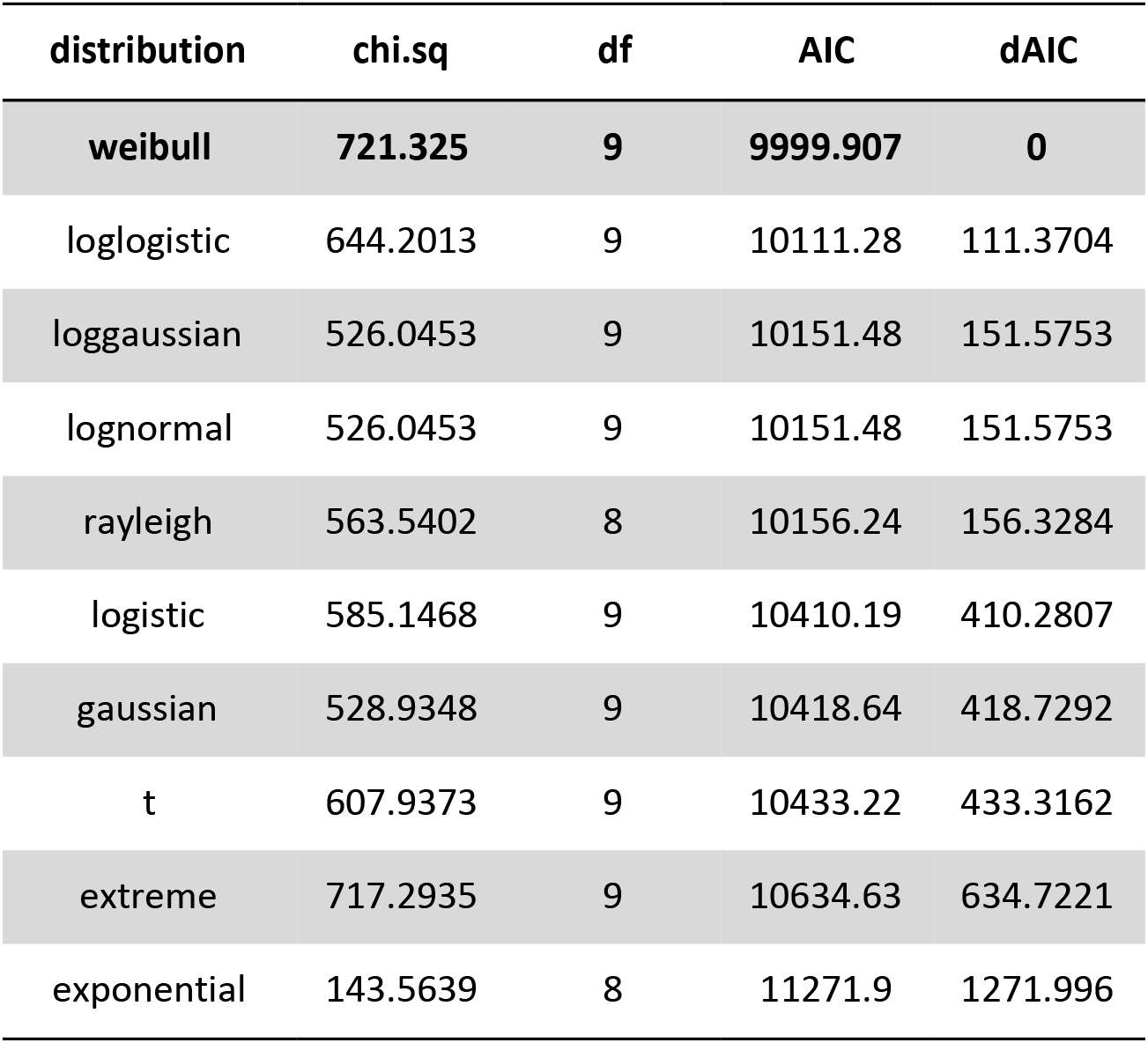
Akaike’s Information Criterion values for distributions tested using accelerated failure time models.

### Wing morphology under tested conditions

As priming began in the larval environment, and larval conditions are known to affect adult morphology [28], we investigated the role to which these environmental conditions modified adult size measured via wing length and area. Priming had a strong effect on wing lengths for both M and S form mosquitoes (2.82 to 3.13mm mean wing length with priming for M-form, 2.87 to 3.06mm for S-form, Additional Figure 3). Increase in wing lengths with priming was significantly different between unprimed and primed mosquitoes for both species via dunn test with adjustment for multiple comparisons (*p*-adj < 0.001).

### Proof of viability post climatic modulation

To evaluate the viability of the longevity-induced state, we moved separate groups of mosquitoes that had lived 66 days in the refugia condition (70 days old) without males to normal insectary conditions (27 °C, 12:12 L:D) and provided an oviposition cup (without blood meal). No eggs were laid in these oviposition cups until the females were allowed to blood feed (day 72 of life) at which point eggs were laid (77 eggs/18 fed female primed S-form, 13 eggs/10 fed female primed M-form). Eggs were hatched and allowed to develop until adulthood. This was successful for primed M and S form mosquitoes (20 emerged S-form adults, 1 emerged M-form adult).

## Discussion

Aestivation appears to be essential to the persistence of mosquitoes in arid, Sahelian regions [29–31], however, we have limited to no understanding of the mechanisms, inductive cues, or prevalence of aestivation in *Anopheles* species [19]. A primary reason behind this limited knowledge of aestivation is the current inability to replicate this state under modern laboratory conditions [18, 32]. This is counter to historic studies which successfully induced high longevity in mosquitoes recently collected from the field (156 and 206 days, respectively) [10, 33], but reported limited details on how this induction was achieved. To the best of our knowledge this study is the first of the modern era to extend the maximum lifespan of *Anopheles* mosquitoes over 100 days, with 2.2 to 3.5 fold increase in maximum longevity over standard insectary conditions depending on species. However, we fail to reach the dramatic extension of these historic studies or the 7 fold extension known to be possible in wild *Anopheles* [34]. The life spans present are significantly higher than past studies attempting to induce aestivation states [17–19], however there are no known markers of aestivation in *Anopheles*, thus making the declaration of this state unequivocally as aestivation difficult. Furthermore, the demonstration (though on a small scale) that mosquitoes after 2+ months in refugia, when returned to wet season conditions, will blood feed and lay viable eggs, shows the viability of these potential aestivators to restart a population (even in absence of fresh male mosquitoes).

As we do not know exact refugia conditions in the wild, but know that *Anopheles* mosquitoes prefer cooler, more humid refugia conditions [35], this study uses realistic, but likely somewhat exaggerated, climate conditions to induce aestivation with a reduced temperature and photoperiod. Temperature was the major climatic variable found to extend lifespan in this study, accounting for roughly a doubling of longevity alone (Table 2). The longevity of some dipterans is known to increase below 27 °C (our normal insectary temperature), i.e. *Calliphora stygia* blowflies have an average longevity of 27.7 days at 25 °C, 45.1 days at 20°C, and 90.8 days at 12 °C [36] and *Culex pipiens* female mosquitoes can be kept in a non-diapausing cold storage for up to 102 days at 6 °C [37]. However, *An. gambiae* s.s. lifespans have been reported largely to decrease in longevity below 20 °C with mean survival of 31.77 days for females at 80% RH and maximum overall survival at 15 °C / 100% RH of less than 70 days [38]. Additionally this relationship with temperature appears to be variable, with another study reporting median survival at a similar temperature (21 °C) of 23.2-24.9 days [39]. The closest reference to the over 100 day survival seen in this study was with recently colonized mosquitoes held in dry season semi-field environments (mean temperatures of ~22 °C) in Kenya, with maximum lifespans reaching 92 days [40]. This may indicate the importance of using low generation from the field mosquitoes for future lifespan studies and may also indicate that temperature alone is not the sole driver of long-lived states.

The second most significant variable found in this study was photoperiod. While this variable had previously been explored in the context of *Anopheles* aestivation [18], the light duration was 11 hours/day, rather than 8 hours/day used in this study. In new world *An. quadrimaculatus* and *An. crucians*, a 8:16 L:D cycle induced greater longevity, though this effect was modest (mean life span 25.64 vs. 19.33 days in female *An. quadrimaculatus*, 20.43 vs. 17.24 in female *An. crucians*) [41, 42]. The Senegalese grasshopper (*Oedaleus senegalensis*) collected in Mourdiah, Mali (14.472,−7.473) have shown a strong interaction between photoperiod and temperature in diapause induction in eggs [43], with maximal induction at 25 °C (the lowest temperature tested) and less than 12 hours of light. However, this location has a slightly higher latitude, and thus a greater seasonal shift in daylength, than the original collection location of our M-form strain. Recent work has shown that conditions that induce winter diapause in European *Drosophila* spp. can also induce the state in some African/tropical strains of flies [44, 45]. This suggests that diapause may be a generalized survival mechanism that is often preserved in tropical taxa and can be utilized to bridge the gap between seasonally favorable climatic conditions. This may indicate that by pushing our mosquitoes into exaggerated, but not necessarily impossible, temperature/light refugia conditions we increased our chances of entering a long-lived state. This exaggeration may be necessary as laboratory conditions do not recapitulate all of the possible cues present in the field. This may explain why our S-form, *An. gambiae* s.s., mosquitoes which were colonized from Cameroon appear to have a similar response to these cues though they are not believed to be aestivating in Mali [12]. This may point to why the interaction term in our model failed to show an effect with temperature or light with species (Table 2).

Consistent with previous studies investigating possible dry-season / aestivation-induction conditions [17, 18], wing sizes were significantly increased in our dry-season priming conditions (Additional File 4). However, there was not a consistent correlation between the wing size and length of life (Additional File 5). As wing size (a good proxy for overall body size) of mosquitoes is primarily driven by larval environments [46], and aestivating mosquitoes are thought to be surviving as adults [12, 32, 34, 47], the exact role of body size in aestivation is unknown, though seasonal differences have been reported in the field [48]. The role of nutrition, long known to play a role in longevity [19, 49–52], may also be driving some of the morphological associations seen in this state. Additionally nutrition may be modulated by how temperature is affecting the overall activity of the mosquitoes. A reduction in activity could also have a reduction in sugar feeding, which may contribute to the increase in longevity seen.

**Additional File 4:**
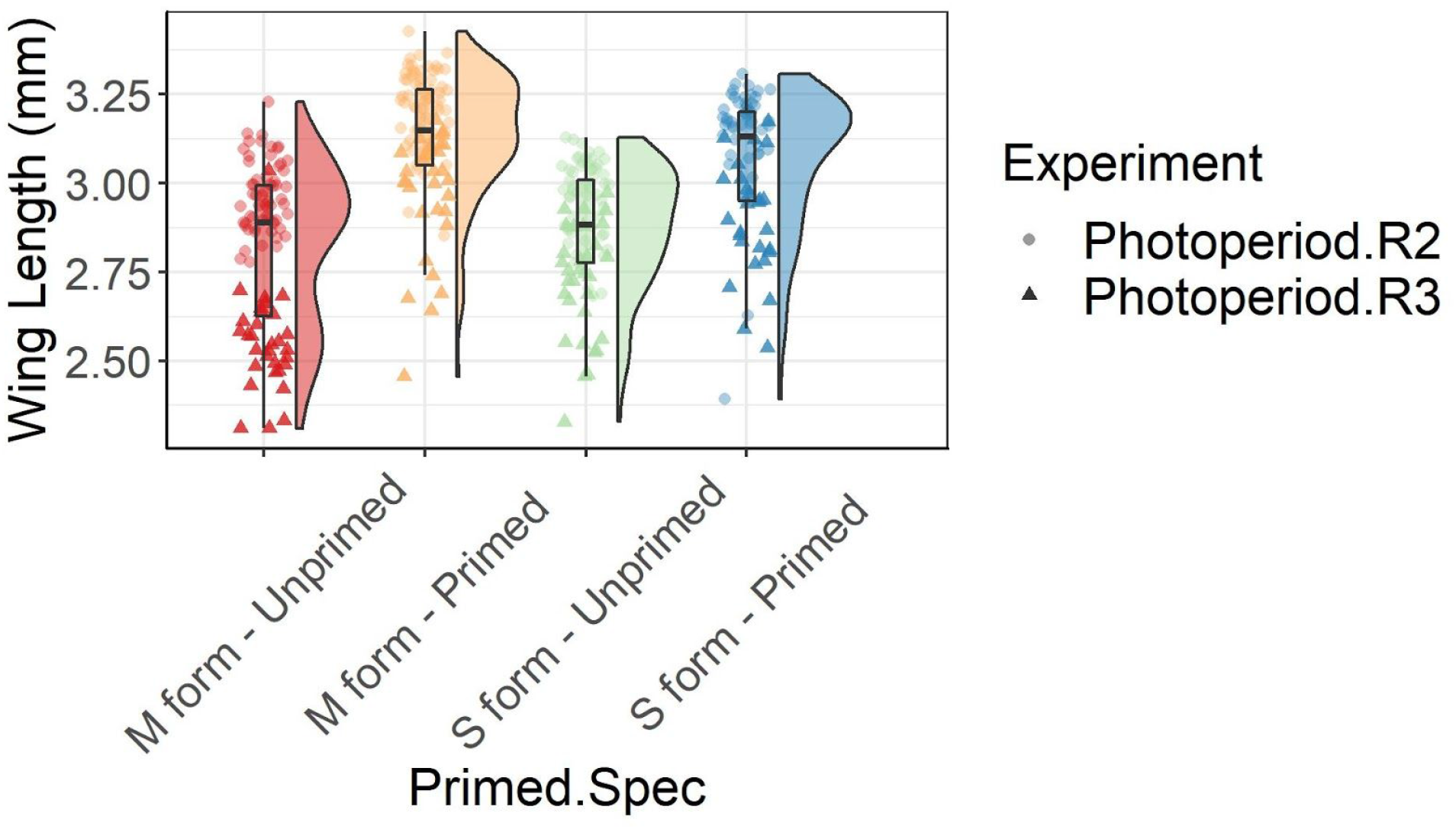
Raincloud plot indicating measured wing lengths as a proxy for body size for primed and unprimed mosquitoes. Boxplot, individual points, and density distribution are shown. Point shapes indicate which replicate the lengths are from.

**Additional File 5:**
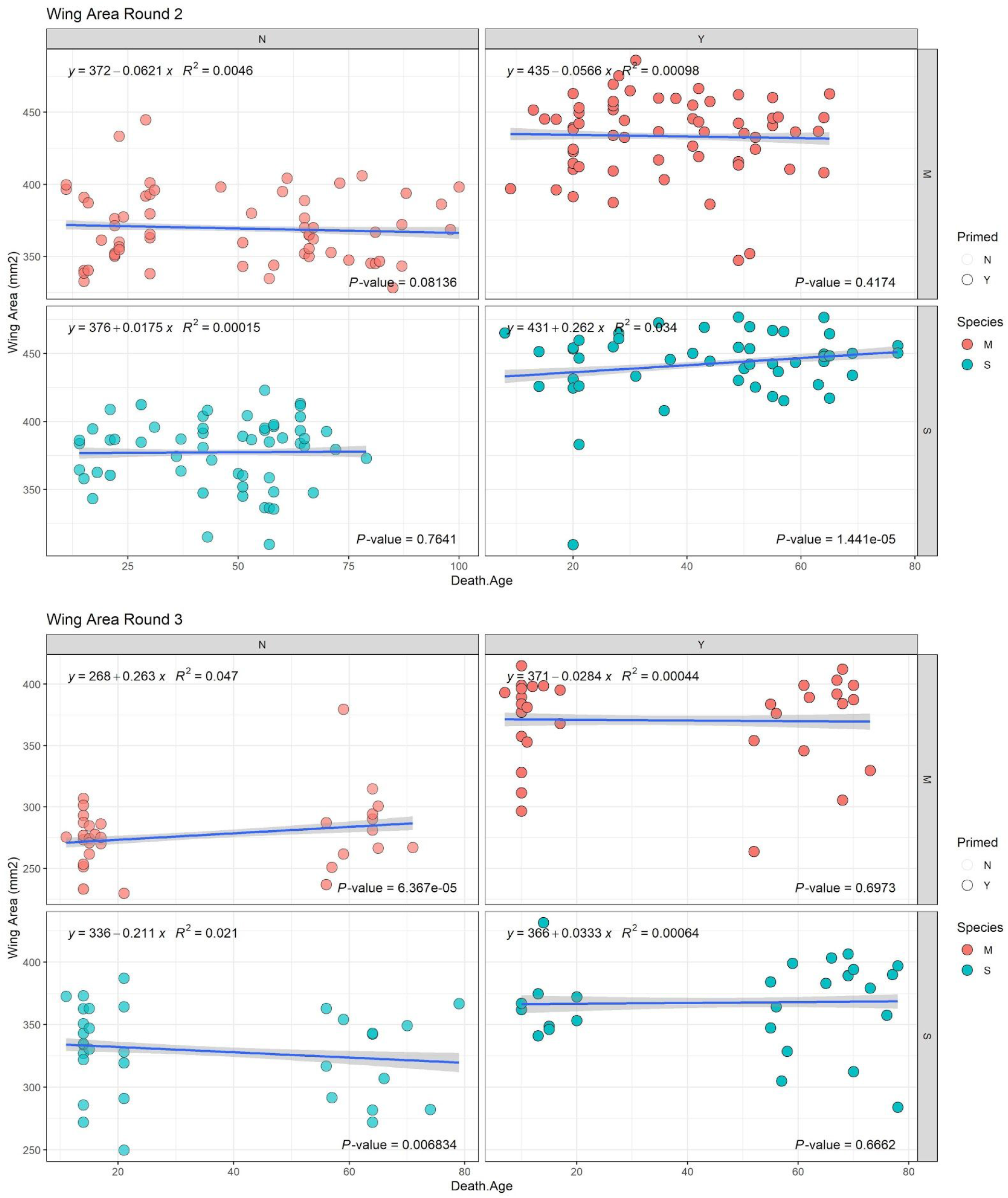
Correlations of wing areas and survival for replicates 2 and 3 of the experiment. No consistent correlations between wing area and lifespan were found between the two experiments.

## Conclusions

This work represents a significant increase over past attempts to induce aestivation in laboratory *Anopheles gambiae* s.l. mosquitoes, possibly representing the first modern recapitulation of the state in the laboratory. Additional fine tuning of priming and maintenance conditions may further push longevity. Future work based on the treatments described here is necessary to validate the state induced in mosquitoes in these conditions through analysis of transcriptional changes [15, 53, 54], ovary follicle length [55–57], lipid content [55], and sugar utilization [58]. These should be contrasted with field-collected samples which may better represent the full expression of the aestivation phenotype. Finally, this work may provide a platform to assess how longevity and aestivation affects *Plasmodium* competence and the possibility of a parasite reservoir in aestivating mosquitoes which may impact future elimination campaigns.

## List of Abbreviations

RH: Relative Humidity
L:D: daily light to dark hours
M-form: *Anopheles coluzzii*
S-form: *Anopheles gambiae* s.s.
AIC: Akaike Information Criterion

## Declarations

### Ethics approval and consent to participate

Not applicable.

### Consent for publication

All authors have reviewed and consented to the publication of this work

### Availability of data and materials

“The datasets supporting the conclusions of this article are available in the zenodo and github repositories, at: zenodo: 10.5281/zenodo.3735020 and https://github.com/benkraj/aestivation.manu.files.scripts

### Authors’ contributions

BK RF TL designed the study. MS dissected and digitized wings. BK LV LG took daily survival data. BK TL analyzed the data. BK TL wrote the manuscript. All authors edited the manuscript.

## Acknowledgements

We thank BEI resources and the original depositor for the N’dokayo strain of *An. gambiae* s.s. mosquitoes used in this study.

